# Sexual isolation with and without ecological isolation in marine isopods *J. albifrons* and *J. praehirsuta*

**DOI:** 10.1101/260489

**Authors:** Ambre Ribardière, Elsa Pabion, Jérôme Coudret, Claire Daguin-Thiébaut, Céline Houbin, Stéphane Loisel, Sébastien Henry, Thomas Broquet

## Abstract

Sexual barriers associated with mate choice are nearly always found to be associated with some level of ecological isolation between species. The independence and relative strength of sexual isolation are thus difficult to assess. Here we take advantage of a pair of isopod species (*Jaera albifrons* and *J. praehirsuta*) that show sexual isolation and coexist in populations where they share the same microhabitat or not (i.e. without or with ecological isolation). Using no-choice trials and a free-choice experimental population, we estimated the strength of sexual isolation between *J. albifrons* and *J. praehirsuta* individuals originating from these different ecological contexts. We found that sexual isolation is strong in presence and absence of ecological isolation, but that it is asymmetric and fails to prevent gene flow entirely. First-generation post-zygotic barriers were low, and there was no sexual isolation within *J. praehirsuta* across habitats. The *J. albifrons* / *J. praehirsuta* species pair thus provides an example where the role of sexual isolation as a barrier to gene flow i) does not depend upon current ecological isolation, ii) seems to have evolved independently of local ecological conditions, but iii) is insufficient to complete speciation entirely on its own.

## Introduction

Sexual isolation resulting from divergence in mating choice is common between closely related animal species, and the evolution of this type of barrier is considered to be a major component of speciation (Coyne & Orr, 2004). However, sexual barrier effects are often found in conjunction with some level of ecological isolation, raising the following questions (discussed e.g. in Ritchie, 2007, Maan & Seehausen, 2011): What are the current relative contributions of sexual and ecological barrier effects in reproductive isolation between animal species? How often, if ever, has sexual isolation initiated speciation rather than evolving secondarily to ecological isolation? And may sexual isolation be an independent driver of speciation or is the evolution of sexual barriers necessarily linked with that of ecological isolation? Similar questions extend to other isolating barriers as well, but we focus here on the relationship between sexual and ecological isolation, and more precisely on the last question, pertaining to the interdependence of these two isolating barriers.

Sexual isolation may be tightly linked with ecological isolation for several reasons (see Butlin & Smadja, 2018 for a discussion of coupling mechanisms and their importance in speciation). A direct form of coupling happens when some genes or traits affect ecological and sexual barriers at once (reviewed in Servedio et al., 2011). In *Heliconius* butterflies, for example, wing colour patterns affect both mimicry and mating signals, resulting in a strong combination of ecological and sexual barrier effects (Jiggins et al., 2001). Along the same line, sexual and ecological barriers will coevolve when they involve overlapping metabolism networks, or sets of genes that are physically linked on the genome. Furthermore, when there is no such intrinsic interdependence between barrier effects, other forms of coupling can happen if environmental conditions promote ecological isolation and simultaneously affect sexual isolation mechanisms. This effect can be particularly strong when natural selection is involved in some aspect of the sexual barrier. Such situations occur when the environment has an effect on male phenotypes, mortality costs associated with sexual display or choosiness, or the transmission of sexual signals (e.g host-dependent sexual signalling in *Enchenopa* treehoppers McNett & Cocroft, 2008). All these cases may lead to the simultaneous evolution of sexual and ecological isolation (reviewed in Maan & Seehausen, 2011, Nosil, 2012, Safran et al., 2013, Boughman & Svanback, 2017, Servedio & Boughman, 2017). This list should even be extended if one considers not only the behavioural aspects of sexual isolation but also gametic isolation. The impact of ecological differentiation in fact appears so ubiquitous that one can wonder in what conditions may sexual isolation ever evolve independently from ecological isolation.

Sexual selection mechanisms such as the Fisher-Lande process of coevolution between arbitrary male traits and female preferences can theoretically drive reproductive isolation largely independently of environmental heterogeneity and ecological barrier effects. This is also perhaps possible in some cases when good genes or compatible genes systems drive sexual isolation between populations. An objective of empirical research is thus to explore how sexual isolation mechanisms are connected to ecological conditions and preferences, and evaluate to what extent sexual isolation may act as an independent driving force in speciation.

Hybrid zones provide good opportunities to investigate the interplay between different types of isolating barriers, including sexual and ecological isolation. Reviewing hybrid zone case studies with contrasted levels of admixture, Jiggins and Mallet (2000) have highlighted that hybridizing animal species where parental genomes maintain a high level of cohesion and most individuals resemble the parental forms (that is, bimodality) are characterized by a strong level of sexual isolation. But the authors also suggested that habitat-mediated exogenous selection is required to maintain the stability of such bimodal hybrid zones. More generally, there are in the literature many more hybrid zone case studies reporting sexual isolation in conjunction with ecological isolation rather than cases where sexual isolation appears to be the most important isolating barrier with little or no ecological isolation. The European hybrid zone between carion and hooded crows is one rare example of a situation where a (slight) level of isolation is maintained by sexual barriers only (Haas et al., 2010, Poelstra et al., 2014). Other examples come from situations where sexual isolation is reinforced by selection against hybridization. For instance, *Mus musculus musculus* and *M. m. domesticus* subspecies of the house mouse are partially isolated by sexual barriers and this has nothing to do with ecological factors (Smadja et al., 2004, Smadja & Ganem, 2005).

The most detailed information should come from situations where one can investigate the role of sexual isolation in different ecological contexts. This is possible in mosaic or otherwise replicate hybrid zones (reviewed in Harrison & Larson, 2016) where hybridizing taxa meet repeatedly in different locations (or different transects can be analysed within a large hybrid zone). In such studies, even when sexual isolation was proven to be a critical barrier to gene flow between species, it appeared to be nonetheless strongly dependent upon ecological conditions. As mentioned above, this happens when sexual and ecological isolation involve the same traits or sexual isolation mechanisms are linked with ecological conditions. Habitat heterogeneity may then have driven ecological isolation and sexual isolation simultaneously (e.g. speciation in *Gasterosteus* sticklebacks and *Pundamilia* cichlids driven by adaptation of female perceptual sensitivity to ambient light combined with sexual selection on male colour, Boughman, 2001, 2002, Seehausen et al., 2008). With these examples the important point is that, whatever its strength, sexual isolation is likely to break down when ecological conditions are homogenized (Seehausen, 2009, see also Taylor et al., 2006). There are also many other cases of mosaic or replicate hybrid zones where sexual isolation is not fully understood but where ecological isolation or spatial segregation, regardless of other isolating barriers, appeared fundamental to the maintenance of reproductive isolation (e.g. field crickets, Harrison & Rand, 1989, marine mussels, Bierne et al., 2003, swordtail fish, Culumber et al., 2011, river herrings, Hasselman et al., 2014, lampreys, Rougemont et al., 2015).

There are comparatively few cases of replicated hybrid zones where sexual isolation appears to be strong and essentially independent of habitat heterogeneity, and a fortiori, independent of ecological isolation. A potential example is the hybrid zone between *Chorthippus* grasshoppers in northern Spain, where female mate choice based on male calling songs generates strong premating isolation that seems not tightly linked with ecological differentiation (Bridle et al., 2001, Bridle et al., 2002, Bailey et al., 2004, Bridle et al., 2006). Another example is the European house mouse hybrid zone, already mentioned above. This system has been studied repeatedly in distant regions, confirming the role of sexual isolation regardless of geographic and ecological conditions (Smadja et al., 2004, Bimova et al., 2011).

When sexual isolation is found in conjunction with ecological isolation, it is interesting to understand the relative roles and interdependence between these two types of barriers. It informs us on the mechanisms that are currently shaping species boundaries, and in some cases on the origin and evolution of these mechanisms (Boughman, 2001, Jiggins et al., 2001, Seehausen et al., 2008).

Here we focus on isopods *Jaera albifrons* and *J. praehirsuta*, two closely related species that show strong sexual isolation and generally occupy distinct habitats but can also be found in a region where they coexist in the same habitat, therefore allowing us to ask whether sexual isolation stands in a situation where ecological isolation doesn’t.

The two species *J. albifrons* and *J. praehirsuta* both belong to the *Jaera albifrons* complex. This complex is composed by five species of small (2-5 mm) marine isopods that live on the shores of the temperate and cold waters on both sides of the North-Atlantic Ocean (Bocquet, 1953, Solignac, 1978). All species of the complex are phenotypically indistinguishable except for male secondary sexual traits that are used for tactile courtship (Bocquet, 1953, Solignac, 1981). The males of each species have specific sets of setae and spines located at different places on their peraeopods (Fig. 1) and they use these features to brush a particular region of the back of females in order to get them to engage in sexual intercourse. The males of both species share the same basic courtship behaviour, whereby they mount females in a head-to-tail position and exercise their brushes. Females accept or reject a male based on this tactile stimulus, and behavioural isolation is thought to ensure a nearly complete arrest of interspecific gene flow in nature (Solignac, 1978).

**Figure 1.**
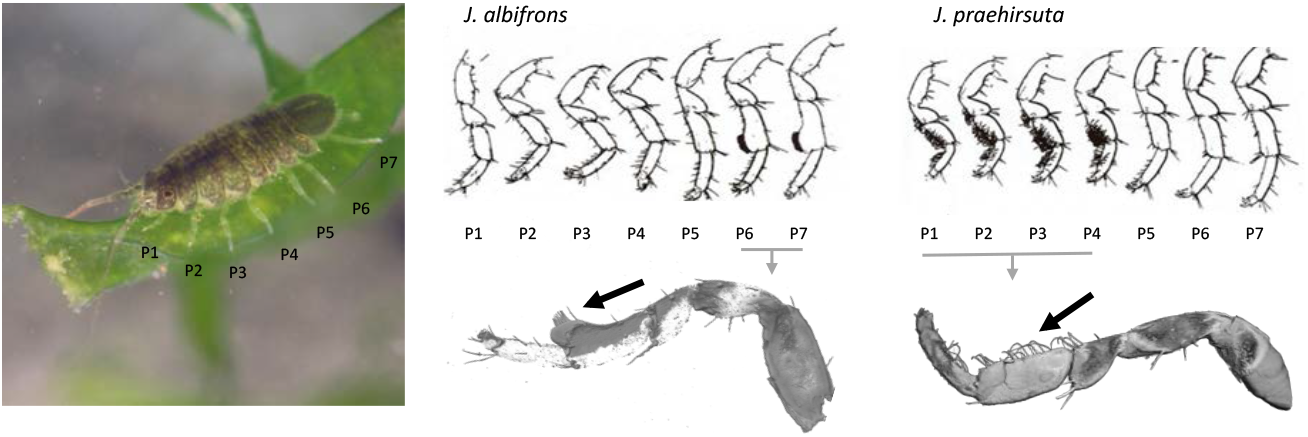
Sexual traits used for tactile courtship by male *Jaera albifrons* and *J. praehirsuta*. In male *J. albifrons*, the second segment (carpus) of peraeopods 6 and 7 extends as a lobe bearing a patch of setae (indicated by the black arrow on the left). Male *J. praehirsuta* instead have curved setae distributed on the first three segments (propus, carpus, merus) of peraeopods 1-4 (right black arrow), and one or two spines on the carpus of peraeopods 6 and 7. The photo on the left shows a 4mm-long adult female (photo credit to Guillaume Evanno & Thomas Broquet). After fluorescent labelling of dissected appendices, close-up pictures were obtained with a confocal laser scanning microscope and processed using software Fiji and IMARIS (photo credit to Sébastien Colin).

Sexual isolation is thus currently a very important barrier between these species, and it could have initiated speciation (Solignac, 1981). However, the five species of the complex also show some level of habitat segregation according to position on the shore, exposure, salinity, and substrate (Naylor & Haahtela, 1966, Jones, 1972). Most remarkably, while *J. albifrons* and *J. praehirsuta* occupy the same narrow belt of intertidal habitats along the American and European shores of the North-Atlantic Ocean, *J. albifrons* is primarily found under pebbles and stones while *J. praehirsuta* is primarily found on intertidal brown algae (at least along European coasts, Bocquet, 1953, Naylor & Haahtela, 1966, Naylor & Haahtela, 1967). These habitats are often in immediate proximity, and these preferences are not strict (Solignac, 1981, Ribardière et al., 2017), but they still imply that ecological isolation is strong and thus its current relative contribution to total reproductive isolation must be important (because this is the first barrier to occur).

The *J. albifrons* / *J. praehirsuta* pair gives us an opportunity to examine the relative strength and interdependence of sexual isolation and ecological isolation because these two species that usually use distinct habitats were reported to coexist in an exceptional population where they share the same habitat (under stones, that is, the primary *J. albifrons* habitat, Solignac, 1969b, a). This coexistence of the two species in a unique habitat was recently found to have persisted for decades and to be more widespread than previously thought as it extends to several other sites at least in the French region Normandy and in the United Kingdom (Ribardière et al., 2017; see also Mifsud 2011). Hybridization happens in these populations (Solignac, 1969a) and results in various levels of introgression (Ribardière, 2017, Ribardière et al., 2017), pointing toward reduced reproductive isolation. Interestingly however, in these hybridizing populations most males bear sexual traits that are clearly identified as belonging to one or the other species and intermediate phenotypes are scarce, suggesting that reproductive isolation does not break down completely.

Ribardière et al. (2017) suggested that sexual isolation is one of the components allowing the persistence of bimodality in spite of introgressive hybridization. If this hypothesis is correct and sexual isolation does not disappear in absence of ecological isolation, then it would suggest that sexual isolation has evolved without a direct dependence on ecological conditions and ecological isolation.

Our main objective was to test whether sexual isolation stands in spite of introgressive hybridization in populations showing no ecological isolation. To reach this objective we quantified sexual isolation between *J. albifrons* and *J. praehirsuta* using experimental “no-choice” crosses between individuals that originated either from a region where ecological isolation is strong or a region where ecological isolation is lacking and the two taxa hybridize. Using individuals from this second region we also quantified sexual isolation in a “free-choice” experimental population where females can escape males and there is competition between individuals, unlike in no-choice crosses where mate rejection may be more constrained (e.g. Jiggins et al., 2001).

In addition, no-choice crosses were also performed with individuals from across our two regions in order to test whether sexual isolation could have evolved differently in different ecological contexts. In particular, species *J. praehirsuta* is found on markedly distinct substrates (seaweeds vs pebbles) in our two study areas, giving us the opportunity to test for an effect of this ecological difference on sexual isolation between populations.

Finally, we took advantage of our experimental crosses to check for potential first-generation post-zygotic barrier effects. While post-zygotic isolating barriers are more likely to operate at later generations, chromosomal differences have been reported in our two focal species (Staiger & Bocquet, 1956, Lécher & Prunus, 1971). Thus we took the opportunity of our experiments to check for the possibility that such differences have an effect already from the first generation of hybridization.

## Methods

### Species

Contrary to males, females of the five species within the *Jaera albifrons* complex are morphologically indistinguishable. They follow the same reproductive cycle (total duration ca. 3 weeks) during which embryos develop in a marsupium (brood pouch) for about 12 days (Solignac, 1976). Development is direct, there is no pelagic larval stage, and offspring measure ca. 0.5 mm when they are released from the marsupium. Individuals become sexually mature and can be sexed within 4 to 5 weeks based on praeoperculum differentiation (e.g. Solignac, 1979).

### Experimental set-up

Our study is based on the analysis of the reproductive output of virgin males and females used in intra- and inter-specific controlled mating experiments (set-up detailed in Fig. 2). In theory, whether or not juveniles are produced in these experiments could result not only from sexual barrier effects but also post-mating pre-zygotic or post-zygotic barrier effects (e.g. inviability of hybrid embryos). However, all past analyses of inter-specific crosses in the *Jaera albifrons* complex have shown that females either rejected hetero-specific males or produced a normal number of offspring. That is, females that produced no offspring did not mate (e.g. Solignac, 1978 p. 49), and females mated by a heterospecific male did not show any reduction in fecundity (e.g. Solignac, 1978 pp. 80-82). There is no postmating copulatory behavioural isolation or mechanical isolation (Bocquet, 1953 p. 297, Jones & Fordy, 1971, Veuille, 1978). The complete absence of offspring produced by a pair of individuals is thus a good indicator for sexual isolation (and most probably the behavioural component of sexual isolation, although gametic isolation has yet to be investigated in this group, see discussion).

**Figure 2.**
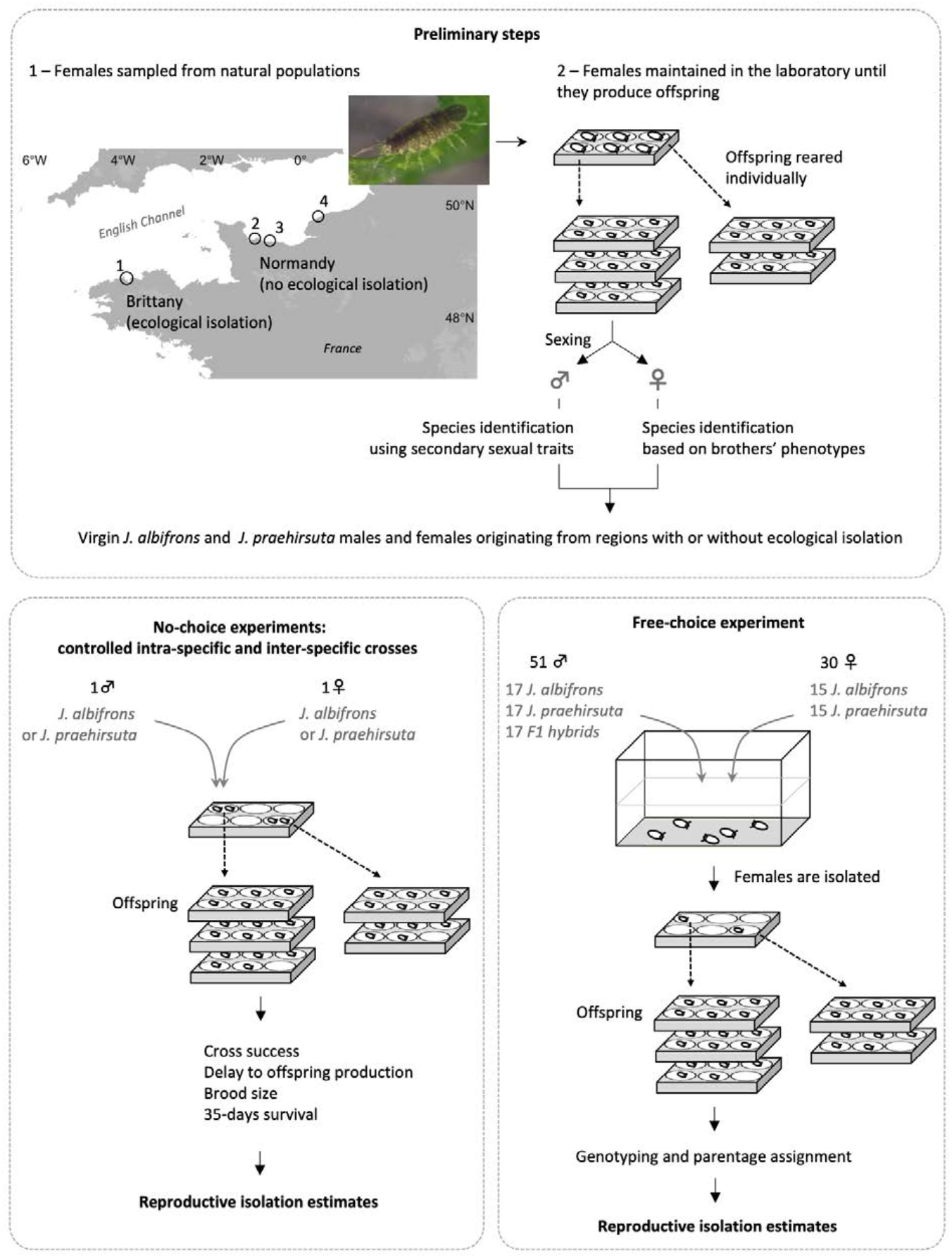
Outline of experimental protocols. The upper panel presents the preliminary steps that were taken to obtain virgin males and females of each species, which could then be used in controlled experiments. Adult females were sampled from natural populations as shown on the map. We knew from previous genetic analyses that local sympatric populations of *J. albifrons* and *J. praehirsuta* were reproductively isolated (region Brittany) or not (introgressive hybridization, region Normandy). Adult females were kept in the laboratory until they produced offspring that were then raised until the male offspring could be identified. Virgin individuals born in the lab were then chosen to be used in no-choice cross experiments or a free-choice experimental population as described in the main text and supplementary information.

To obtain virgin individuals of both sexes and both species, we first sampled (unidentified) females in natural populations where *Jaera albifrons* and *J. praehirsuta* coexist (Fig. 2). We chose populations where we knew from previous work (Solignac, 1978, Ribardière, 2017, Ribardière et al., 2017) that the two species leave on different substrates (pebbles *vs* seaweeds, populations from Brittany) or share the same substrate (pebbles, populations from Normandy). Then we individually reared in the lab the offspring produced by these females (which were fertilized by unknown males in nature prior to sampling) until they could be sexed and males could be identified. Females were sorted as *J. albifrons* or *J. praehirsuta* according to the sexual traits held by their brothers. At this stage we thus had a series of *J. albifrons* and *J. praehirsuta* virgin adults originating from populations with or without ecological isolation (region “Brittany” *vs* region “Normandy”). These individuals could then be used in the controlled experiments described below. Female sampling and experimental conditions are detailed in supplementary information.

### No-choice crosses within each region

In order to understand reproductive isolation processes with or without ecological isolation, we first ran a series of crosses where one male and one female from the same region were paired and their reproductive output monitored. These crosses featured intraspecific and interspecific crosses using either a pair of individuals from Brittany (where ecological isolation is strong) or a pair from Normandy (where there is no ecological isolation).

We monitored 23 intraspecific and 17 interspecific crosses within each of these two conditions (that is, 40 crosses within each region of origin; details in Table 1). These numbers were somewhat constrained for three reasons. First, the number of individuals available for the experiment depended on the (unknown) species identity of the females sampled in the wild, their survival and fecundity in the lab, and the survival of their offspring (see preliminary steps in Fig. 2). Second, and more importantly, crosses were designed so that the male and the female that were paired never shared the same mother. Third, each individual was used in a unique cross, so that each cross was an independent replicate.

**Table 1 -.** Number of experimental “no-choice” crosses performed in order to estimate reproductive isolation between marine isopods *J. albifrons* and *J. praehirsuta* in two French regions where they are ecologically isolated (Brittany) or not (Normandy). Intra- and interspecific crosses were performed using males and females from the same region (intra-region) and opposite regions (inter-region).

| Cross type | Female | Male | Intra-region |  | Inter-region |  |
| --- | --- | --- | --- | --- | --- | --- |
|  |  |  | Brittany | Normandy | Britt. fem.<sup>†</sup> | Norm. fem.<sup>‡</sup> |
| Intraspecific |  |  |  |  |  |  |
|  | *J. albifrons* | *J. albifrons* | 17 | 6 | 10 | 10 |
|  | *J. praehirsuta* | *J. praehirsuta* | 6 | 17 | 10 | 10 |
| Interspecific |  |  |  |  |  |  |
|  | *J. albifrons* | *J. praehirsuta* | 10 | 10 | 10 | 10 |
|  | *J. praehirsuta* | *J. albifrons* | 7 | 7 | 10 | 10 |

† Female from Brittany
‡ Female from Normandy

For each cross we recorded i) if it successfully produced offspring, ii) how long it took for the first offspring to appear, iii) how many offspring were contained in each brood produced, and iv) offspring survival at 35 days. These data were used to estimate reproductive isolation components as described below. All analyses were performed in R v.3.3.3 (R Core Team, 2017).

### Reproductive isolation estimated from no-choice crosses within each region

Ecological, sexual, and first-generation post-zygotic components of reproductive isolation were quantified following Sobel and Chen (2014) using estimators that vary between −1 (complete disassortative mating, probability of interspecific gene flow = 1) to 1 (complete reproductive isolation, probability of interspecific gene flow = 0). To compare reproductive isolation components in presence *vs* absence of ecological isolation, all the computations described below were performed independently using crosses featuring individuals “from Brittany” on one hand, and “from Normandy” on the other hand (that is, two independent sets of analyses).

First, because *J. albifrons* and *J. praehirsuta* in Brittany do not have strictly non-overlapping habitats, we used survey data from Ribardière et al. (2017) to quantify ecological isolation as

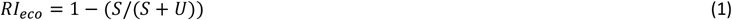

where *U* was the proportion of individuals found on the primary habitat of their species and *S* was the proportion of individuals found on the alternative habitat (i.e. *S* is the probability that an individual is in a place where it will meet the other species, that is, “shared”). This equation gives an estimate of the reduction in interspecific gene flow that would happen if individuals would mate randomly within each habitat (i.e. ecological isolation only).

Second, we estimated three components of sexual and post-zygotic isolation (listed in Table 2) from our experimental crosses. The strength of each reproductive isolation barrier *i* was estimated as

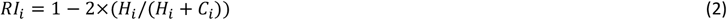

where *H_i_* and *C_i_* refer to variables calculated for heterospecific and conspecific pairs (Sobel & Chen, 2014). For sexual isolation (*RI*_1_), *H* and *C* were the proportions of inter- and intraspecific crosses that successfully produced offspring. For components of post-zygotic isolation, *H* and *C* referred to brood size (number of offspring) or survival (proportion of offspring surviving at day 35) observed from intra- and interspecific crosses (see column “parameter” in Table 2). Each of these components of reproductive isolation was thus estimated independently, giving the strength *RI_i_* that each barrier would have if it were acting alone.

**Table 2 -.**
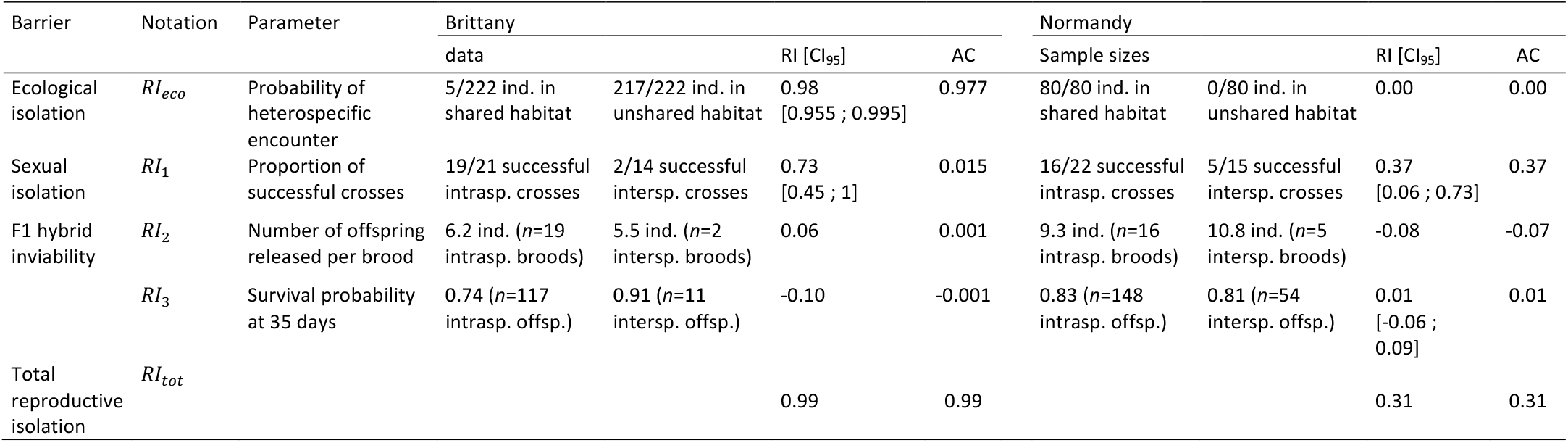
Components of reproductive isolation between marine isopods *Jaera albifrons* and *J. praehirsuta* in two regions with contrasting levels of isolation. Reproductive isolation (RI) was calculated following Sobel & Chen (2014). Bootstrap confidence intervals (CI_95_) based on 10000 resampling of observed data are given whenever sample sizes were not too small (e.g. sample sizes for survival of interspecific offspring in Brittany were particularly small since interspecific crosses were rarely successful). RI gives the limit to interspecific gene flow that would be caused by each specific barrier acting alone. AC gives the actual absolute contribution of each barrier given that other barriers are acting earlier in life cycle. The sum of AC over all barriers is equal to the total strength of reproductive isolation. Sexual isolation contains a strong behavioural component, but could also include a (so far untested) gametic component. RI due to F1 hybrid inviability is based on brood size (number of offspring produced by a mother after intra-marsupial development) and survival of these offspring after 35 days.

A 95% bootstrap confidence interval was calculated for each *RI* estimate by resampling 10000 times the observed data if sample sizes where not too small (i.e. *S* and *U* or *H_i_* and *C_i_* ≥ 14, Table 2).

Total reproductive isolation was estimated using the product of *H_i_* and *C_i_* across all components of isolation (note that we used only multiplicative components of fitness: probability of encounter, probability that a cross is successful, brood size, and probability of offspring survival). Interspecific encounters can only happen in a “shared” ecological context, so *H_i_* values were defined only under the condition that heterospecific individuals meet in nature (and this happens with probability *S*). Conspecific encounters happen in any context (shared or not), but *C_i_* could theoretically take different values (i.e. conditionnal on S and U, Sobel & Chen, 2014). Here because we used no-choice trials, we took *C_i_* values to be equal in any ecological context (e.g. we considered the probability that a conspecific cross was successful to be independent on whether such an encounter would happen in a shared or an unshared context in nature). Total reproductive isolation was therefore calculated as:

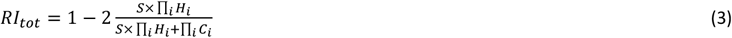

Finally, we estimated the absolute contribution of each individual barrier by subtracting the effect of previously acting barriers as (Sobel & Chen, 2014):

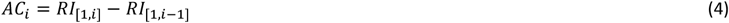

With this definition, individual contributions *AC_i_* can be seen as additive components of *RI_tot_*

### No-choice crosses across regions

In addition to the 80 crosses described so far, we ran another series of 80 intraspecific and interspecific crosses (Table 1) pairing individuals from opposite regions (and thus opposite habitats in the case of *J. praehirsuta*, which rests on algae in Brittany vs. under stones in Normandy). These crosses were useful to test for an effect of habitat on sexual isolation within and between species. They also provided a direct test that *J. albifrons* (or *J. praehirsuta*) from across our two separate regions belong to the same biological species, an implicit assumption of this study and most previous investigations with this system (Solignac 1969b; Ribardière et al. 2017). The reproductive output of these crosses was recorded as described above for no choice crosses within each region.

### Free-choice experiment

The general aim of the free-choice experiment (Fig. 2) was to estimate sexual isolation based on the reproductive output of males and females *J. albifrons* and *J. praehirsuta* freely interacting in an experimental population (and thus experiencing intra-sex competition and easier male avoidance by females, unlike in no-choice experiments). This experiment focused only on the situation where the two species occupy the same habitat in the wild and thus interact frequently. Hence we used virgin adult individuals obtained from the no-choice crosses described above (region Normandy only), so that we could mix *J. albifrons, J. praehirsuta*, and F1 hybrids all obtained in the same controlled conditions and all originating from a region without ecological isolation (Fig. 2). We chose 15 females of each species, 17 males of each species, and 17 males produced by interspecific crosses. These numbers were constrained by several parameters, including the fact that we avoided mixing related males and females (i.e. two males could be brothers, but we did not pick males and females from within the same family). A mixture of 30 virgin females and 51 virgin males of controlled origin therefore composed our experimental population (Fig. 2).

Here we outline the experimental set-up, which is presented in detail in supplementary information. All adults were put together in a small aquarium for 12 days. This is the minimum time required for a female to produce offspring if such a female would have been fertilized early in the experiment (Solignac, 1976). After that, all surviving females were removed from the aquarium and kept individually until they produced offspring, which were then reared individually. All adults were photographed before and after the experiment and genotyped at 13 microsatellite loci (Ribardière et al., 2015). All offspring were also genotyped at the same loci. Photo-identification and genetic parentage assignment (using software Colony v2.0.6.1, Jones & Wang, 2010) were used to identify adult females after the experiment (remember that females of the two species cannot be distinguished otherwise) and identify the father of each offspring.

In addition, the secondary sexual traits of all adult males (Fig. 1) were examined under a microscope to determine their role in male mating success. This is useful in this experiment because free-choice conditions give access to variance in male mating success with females of the two species, and one can thus explore the link between male traits and sexual isolation. Male phenotypes were summarized using principal component analyses (PCA) based upon 13 phenotypic variables (see supplementary information). This approach produces linear combinations of traits that are more efficient for investigating sexual isolation than multiple trait variables separately (Hohenlohe & Arnold, 2010). We used coordinates on the first PCA axis to assess whether female mate choice matched the distribution of male sexual traits within each species (building upon Ryan & Rand, 1993, Arnold et al., 1996).

### Sexual isolation estimated from free-choice experiment

We aimed to compare sexual isolation in these free-choice settings with that measured in no-choice crosses. Hence in a first step we used the same theoretical framework (Sobel & Chen, 2014) to estimate sexual isolation *RI*_1_ following equation (2) with *H*_1_ and *C*_1_ defined as the proportions of inter-specific and intra-specific crosses that successfully produced offspring. These proportions were calculated as the number of successful pairs divided by the number of potential pairs that could possibly have formed. Note that because we initially introduced the same number of males of each species in the experiment, using proportions (as above) or absolute numbers of successful pairs (as in Sobel & Chen, 2014) would lead to the same result for *RI*_1_. It turned out that sexual isolation was very strong: only one interspecific pair and two pairs involving F1 hybrid males produced offspring, while all other successful pairs were conspecific (see results). Hence downstream barriers involving brood size and survival were not quantified because they would be based on too few samples.

In a second step, sexual isolation was estimated using a framework described by Rolan-Alvarez and Caballero (2000) that applies to multiple-choice experiments. While this estimation procedure gives a less direct estimate of interspecific gene flow reduction than Sobel and Chen’s method and cannot be applied to our no-choice experiments, it has several interesting properties. In particular, it takes into account inequalities in mating frequencies rather than assuming that the two species have the same propensity to mate, and it can detect asymmetry in sexual isolation.

To estimate sexual isolation we counted the number of male/female pairs of each type (e.g. *J. albifrons/J. albifrons, J. albifrons/J. praehirsuta*, etc.) that successfully reproduced. Pair sexual isolation (*PSI*) was estimated for every pair type following Rolan-Alvarez and Caballero (2000). This method gives a conservative view of sexual isolation as it is defined for each pair type as the number of observed pairs divided by the number of expected pairs given the actual mating success observed We then followed these authors’ recommendation to estimate *I_PSI_*, a modified joint isolation index (Merrell, 1950) based on *PSI* statistics and that varies from 0 (no isolation) to 1 (complete isolation). Details of the computation are described in Rolan-Alvarez and Caballero (2000) and Perez-Figueroa et al. (2005). Values of *PSI, I_PSI_*, and their statistical significance were computed using JMating v1.0.8 (Carvajal-Rodriguez & Rolan-Alvarez, 2006).

## Results

Out of 160 crosses that were set-up and monitored, offspring were produced in 77 cases. However, a cross was informative only if the female survived for long enough to have a chance to produce offspring. The reproductive cycle of females takes about three weeks (Solignac, 1976), and this figure does not take into account the time needed for mating and the potential delay that depends on the synchronisation between mating and the female’s condition. In this study all but one female that produced offspring survived for at least 24 days after being placed with a male (data not shown). Hence to avoid false negatives we removed from the analyses all crosses (8 intraspecific and 10 interspecific crosses) for which the female survived for less than 24 days after being paired. All the results presented below are thus based on the remaining 142 crosses.

### Reproductive isolation within each region

Estimates of reproductive isolation are presented in Table 2. In region Brittany, Ribardière et al. (2017) reported 141 *J. albifrons* under pebbles *vs* 2 on algae, and 3 *J. praehirsuta* under pebbles *vs* 77 on algae. Ecological isolation can thus be estimated to *RI_eco,Brittany_* = 98%. Because this reproductive barrier is the first to occur in nature, all other barriers will have comparatively little effect in natural populations in this region. This is reflected in the values for absolute contributions (*AC_i_* column in Table 2).

However, as shown in Table 2 and Figure 3, no-choice intraspecific crosses in Brittany were more successful (probability of success=0.9) than interspecific crosses (0.14, Fisher’s exact test p<0.001). These results indicate strong sexual isolation (*Rl*_1,*Brittany*_= 0.73). That is, sexual isolation would reduce interspecific gene flow by about 70% in situations were individuals of the two species would meet, as in the case for instance when bold *J. albifrons* individuals venture on algae, or *J. praehirsuta* on pebbles.

**Figure 3.**
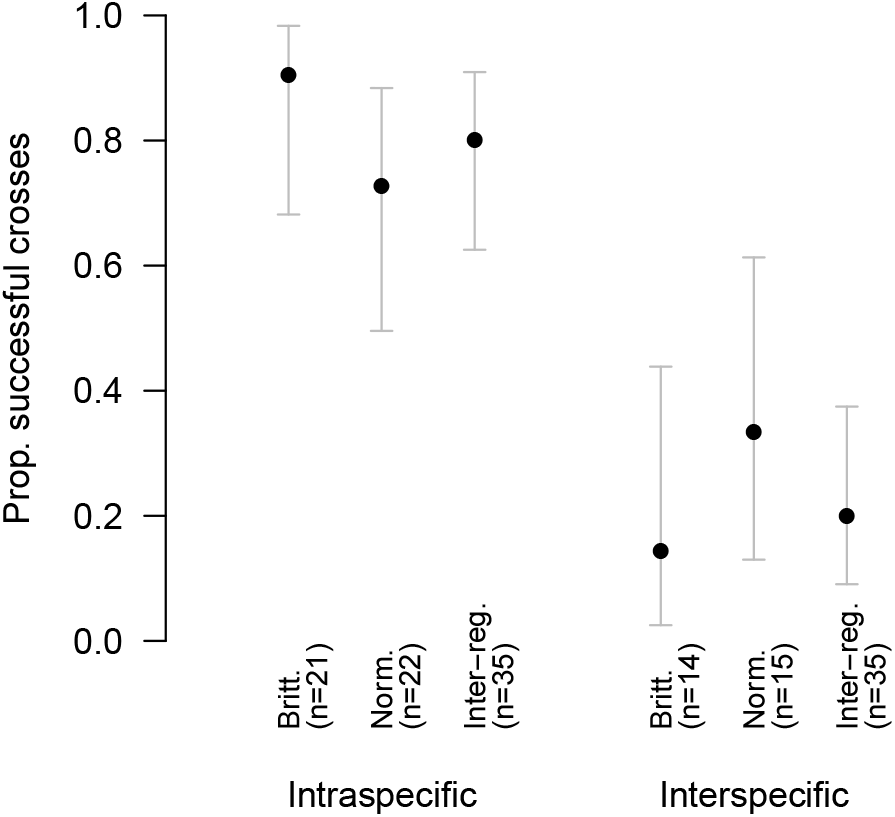
Proportion of successful crosses in no-choice experiments involving intraspecific crosses (either *Jaera albifrons* or *J. praehirsuta*) and inter-specific crosses. The male and female of a given cross could come from the same region (Brittany or Normandy, see text) or each from a different region (inter-reg. crosses). The sample sizes (number of experimental crosses) are given in brackets, and the bars give 95% confidence intervals around each observed proportion. A cross was successful if it produced at least one offspring. Interspecific crosses were consistently less successful than intraspecific ones, pointing towards sexual isolation between *J. albifrons* and *J. praehirsuta*.

By contrast, there was no ecological isolation in mixed *J. albifrons* / *J. praehirsuta* populations from Normandy (Table 2, data from Ribardière et al., 2017) and there we found sexual isolation to be half of that found in Brittany (probability of success = 0.73 vs. 0.33 for intra- and interspecific crosses, *RI*_1,*Normandy*_ = 0.37, Fig. 3). Yet in this region sexual isolation was the first barrier to occur, and thus it effectively led to a strong reduction in interspecific gene flow (*AC*_1,*Normandy*_ = *RI*_1,*Normandy*_ = 0.37). This stands in contrast with the situation described above for Brittany where sexual isolation was twice as strong (*RI*_1,*Brittany*_= 0.73) but had very little effect in nature (*AC*_1,*Britanny*_ = 0.015, Table 2).

The time that a couple of individuals took to produce offspring (Fig. S1) did not enter calculations of reproductive isolation following the framework of Sobel and Chen (2014), but it is interesting to note that intraspecific crosses produced offspring more readily (41.2 days on average, *n*=35) than interspecific crosses (58.9 days, *n*=7, generalized linear model GLM with quasi-Poisson family *p*=0.042). The time needed to produce offspring seemed also more variable among interspecific crosses (Fig. S1). These observations further support sexual isolation in the studied populations (see discussion), but there were too few successful interspecific crosses to look at these data in each region separately.

Successful broods contained from 1 to 33 offspring (mean 7.9 ind.) and on average 79.8 % of the offspring were still alive at day 35. Detailed data for intra- and interspecific crosses within each region are presented in Table 2. These data did not indicate any first-generation post-zygotic barrier effect due to reduced F1 hybrid inviability in either region (Table 2). Accordingly, pooling data from our two regions, brood size (Fig. S2) and survival (Fig. S3) did not differ significantly between intraspecific (*n*=35 broods, mean 7.6 offspring per brood, 79% survival) and interspecific crosses (*n*=7 broods, 9.3 offspring, 83% survival), although there was little power for these tests given that few interspecific crosses produced offspring (brood size: GLM quasi-Poisson family, *p*=0.5, and survival: GLM quasi-binomial family *p*=0.6).

### No-choice crosses across regions

Experiments that crossed individuals from the same species but originating from distinct regions had a high success (28 out of 35 crosses produced offspring, Fig. 3) and this success was similar to that of intraspecific crosses within each region (Fisher’s exact test *p*=0.33). Moreover, this high success was equal for *J. albifrons* pairs and *J. praehirsuta* pairs (respectively 15 out of 19 and 13 out of 16 crosses produced offspring, *p*=1). That is, there was no difference in success when a male and a female came from the same *vs* different regions, and this was true for each of the two species.

Focusing on the differences between intra- and interspecific crosses, all results obtained by crossing individuals from across two distinct regions were identical to the results presented above for crosses within a region. Briefly, intra-specific crosses were more successful (*n*= 35 crosses, probability of success = 0.8) than interspecific crosses (*n*=35, probability of success = 0.2, Fisher’s exact test *p*<0.001, Fig 3), delay to offspring production was shorter in intraspecific (*n*=28 broods, 39.2 days on average) than interspecific crosses (*n*=7, 65.3 days, GLM quasi-Poisson *p*=0.006, Fig. S1), and there was no difference in brood size and survival at day 35 (intraspecific: *n*=219 offspring from 28 broods, 9.21 offspring per brood on average, 81% survival, interspecific: n=68 offspring from 7 broods, 11. 7 offspring per brood, 79% survival, GLM quasi-Poisson and quasi-binomial *p*=0.3 and 0.8, Figs. S2 and S3).

In summary, all differences in reproductive output from intra-versus inter-specific crosses were unaffected by the region of origin of the individuals.

### Results from the free-choice experimental population

We could determine the reproduction patterns for 47 males (out of 51) and 22 females (out of 30), as detailed in the supplementary information. Out of 22 females, most (16) mated with a single male, while 5 had 2 mates and 1 had none (Fig. S4A). By contrast, the majority of males had no success (31 out of 47), while other males mated with 1 to 4 female partners. There were no obvious differences in reproductive success distribution between species (Figs. S5 and S6).

A total of 26 male/female pairs produced offspring. As it turned out (Fig. 4), *J. albifrons* females reproduced only with *J. albifrons* males (*n*=18 pairs), while *J. praehirsuta* females reproduced with the three types of males: *J. praehirsuta* males (*n*=5), “hybrid” males, (*n*=2), and *J. albifrons* males (*n*=1). This distribution is not random (Fisher’s exact test for independence between male and female species composing these pairs p<0.001) and leads to an estimate for sexual isolation between *J. albifrons* and *J. praehirsuta* equal to *RI*_1,*exp.pop*._=0.92 (CI_95_ [0.75;1]), a number that can be directly compared with *RI*_1,*Normandy*_=0.37 obtained from no-choice crosses.

**Figure 4.**
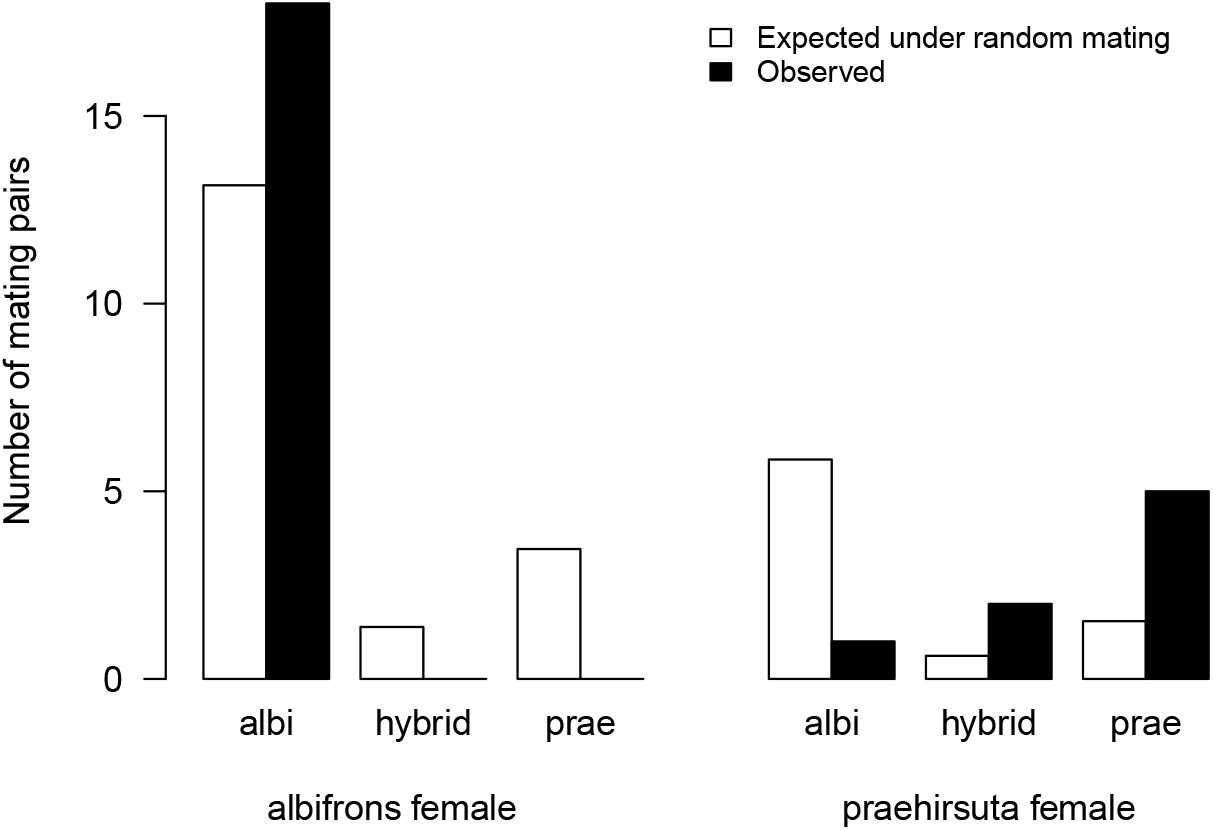
Number of mating pairs of each possible type expected from random mating in absence of sexual isolation (white bars) and observed from an experimental population (black bars). The expected numbers take into account not only the number of males and females of each type that entered the experiment but also their propensity to mate (see text). We see that *J. albifrons* females mated successfully only with *J. albifrons* males, while *J. praehirsuta* females produced offspring with males of the three different types, although not in equal proportions. These results point to strong but imperfect, asymmetrical sexual isolation.

These results assume that all individuals had the same probability to mate. Rolan-Alvarez and Caballero’s method based on the calculation of pair sexual isolation (*PSI*) is more conservative, since it takes into account the effect of differences in mating success between individuals from each species (e.g. here *J. albifrons* individuals mated more frequently than others). Figure 4 shows the number of mating pairs of each possible type expected from random mating in absence of sexual isolation but taking into account the variation in mating frequencies. Estimates of *PSI* are given in Table S1. The global modified joint isolation index *I_PSI_* (Rolan-Alvarez & Caballero, 2000) was equal to 0.46, while pair-specific values were 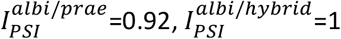, and 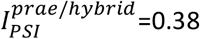, indicating strong isolation between *J. albifrons* and *J. praehirsuta*, and lower isolation between *J. praehirsuta* and hybrids. Bootstrap-based tests performed in JMating for all *PSI* and *I*_PSI_ estimates showed that only 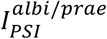 was statistically significant (*p*=0.004).

Most of the variation in parent phenotypes (male secondary sexual traits) could be reduced to two PCA axes (61.8% and 18.3% of the variation explained, Fig. 5A). Using PCA axis 1 to summarize male sexual traits in univariate space, we see in Figure 5B that the preferred mates of *J. albifrons* females coincide with the distribution of *J. albifrons* male sexual traits, while *J. praehirsuta* females mated with males showing a wider range of trait values.

**Figure 5.**
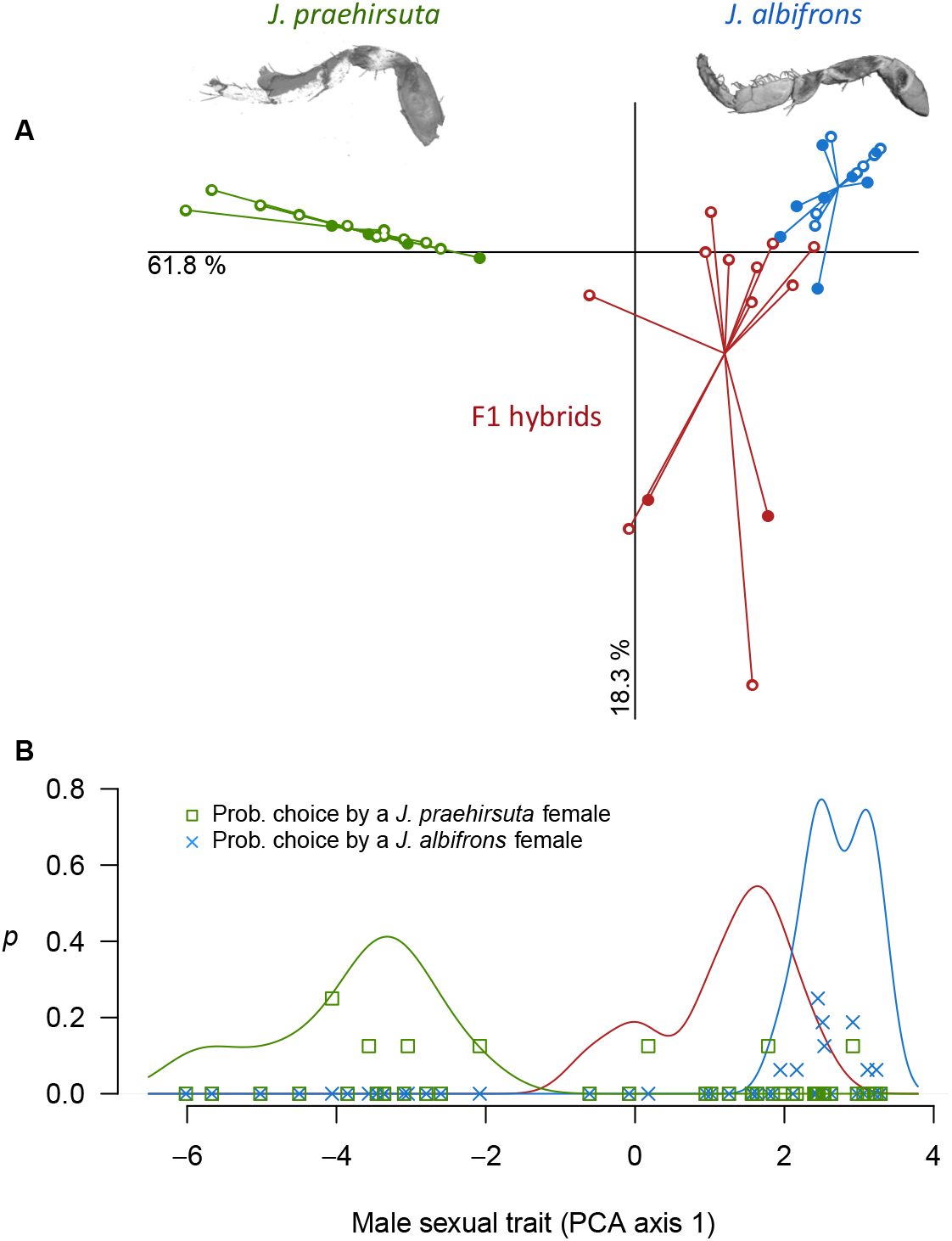
Asymmetric sexual isolation between *Jaera albifrons* and *J. praehirsuta*. Panel A) shows the first two components of a principal component analysis of male phenotypes (potential parents in the free-choice experimental population) grouped by types. We see that individuals of the *albifrons* or *praehirsuta* types are phenotypically differentiated, while individuals of the hybrid type (i.e. produced by an “interspecific” cross) show more phenotypic variability, including phenotypes indistinguishable from the parental morphs. In this PCA plot, empty circles represent the males that did not sire any offspring, while solid dots show the males that successfully reproduced. In panel B), the solid curves show the density distribution of male sexual trait values in univariate space (PCA axis 1 only). The probability that a female mated with a male showing a particular sexual trait value is shown by green squares (*J. praehirsuta* females) and blue crosses (*J. albifrons* females). The preferred mates of *J. albifrons* females coincide with the distribution of *J. albifrons* male sexual trait. By contrast, this concordance is much relaxed for *J. praehirsuta* females, which mated with males showing a wide range of trait values.

## Discussion

In this study we evaluated sexual isolation between *Jaera* species in two different contexts of ecological isolation in order to determine whether sexual barrier effects depend on ecological isolation. For that, we took offspring production in intra-*vs* inter-specific crosses as an indicator of sexual isolation. This is a valid choice because brood size did not differ between intraspecific and interspecific crosses, and the abortion of an entire brood of embryos was never observed (oocytes and developing embryos are visible in the marsupium of females). Hence, conform to what had been reported previously (Solignac, 1978), females in presence of a heterospecific male either produced no offspring because they did not mate, or produced a normal number of offspring. The success of crosses (presence vs absence of juveniles produced) is thus a good indicator of sexual isolation.

### Sexual isolation was effective both with and without ecological isolation

We found strong sexual isolation both in a context where ecological isolation is nearly complete (*RI_eco, Brittany_*=98%, *RI*_1,*Brittany*_ = 73%) and in a context where there is no ecological isolation and individuals of the two species may hybridize (*RI_eco, Normandy_*=0, *RI*_1,*Normandy*_ = 37%). These RI values based on reproductive success in no-choice conditions suggest that sexual isolation is less effective when the two species co-occur in the same habitat (Normandy). However, the free-choice experiment showed that when the conditions allow females to escape or choose amongst several males, then sexual isolation is in fact very efficient even in the hybridizing populations from Normandy (*RI*_1,*exp.pop*._ = 92%).

Moreover, most of our controlled no-choice crosses were monitored for a long time (5 to 196 days, median 22 days), meaning that females were stuck with a given male in a tiny area for weeks and weeks, and thus our estimates probably give us a lower bound on sexual isolation. We conclude that sexual isolation remains strong in populations where the two species share the same habitat, (despite hybridization and introgression, Ribardière, 2017, Ribardière et al., 2017) and thus sexual isolation in this system is largely independent of ecological isolation.

In addition, while males were identified as *J. albifrons* or *J. praehirsuta* based on phenotypes that are directly relevant to reproductive isolation, this was not the case for females, which were identified based on the phenotype of their brothers (see methods, Fig. 2, and supplementary material). Hence in all our experiments we had no direct information on the sexual phenotype of females (preferences). This uncertainty on what really was a “*J. albifrons*” or a “*J. praehirsuta*” female in our experiments makes it all the more remarkable that sexual isolation was found to be strong, especially in the populations where the two species share the same habitat and hybridize.

### Did sexual isolation evolve independently of ecological contexts ?

*J. praehirsuta* individuals in Brittany vs. Normandy dwell on different habitats (pebbles vs. brown algae). But intraspecific crosses had exactly the same success (and the same delay to offspring production) whether or not males and females originated from the same region or different regions (Figs. 3 and S1). And this success was the same as that of *J. albifrons* pairs (within or across regions). Hence habitat differences did not generate any trace of sexual isolation between individuals originating from algae vs. rock populations in species *J. praehirsuta*. Sexual isolation was unaffected by the habitat of origin and thus we can conclude that it did not evolve through context-dependent sexual selection mechanisms that would have selected for divergent, locally adapted, trait-preference regimes.

However, to suit our purpose of estimating reproductive isolation in different populations, all individuals were reared in identical, artificial, lab conditions (see supplementary information) where all five species of the *Jaera albifrons* complex had been shown to have high fitness (Bocquet 1953; Solignac 1978). This means in particular that all individuals used in cross experiments were born and raised in the exact same conditions, regardless of their population of origin. Hence we cannot exclude that contrasted substrates (algae vs. rocks) could have some proximate effect on mate choice mechanisms, for instance through phenotypic plasticity in epicuticular hydrocarbons or other contact chemical cues involved in tactile courtship (e.g. Stennett and Etges 1997). Such chemical signalling mechanisms have not yet been investigated in the *Jaera albifrons* complex.

In addition, as mentioned above we observed from no-choice trials that sexual isolation was twice as strong in populations where the two species are ecologically separated (*RI*_1,*Brittany*_= 73%, *RI*_1,*Normandy*_ = 37%). One plausible explanation is that mate choice is relaxed in introgressed populations, but only in situations where mate options are scarce (in agreement with our finding that sexual isolation was stronger in the free-choice experimental population *RI*_1,*exp.pop*._ = 92%). Variations in the relative abundance and spatial distribution of individuals in natural settings may thus modulate the likelihood of interspecific mating. It is often argued that this could happen when dynamic habitats modulate the intensity of sexual isolation by limiting mate choice options, perhaps in some cases approaching no-choice conditions, if for example a female of species A finds herself isolated for some time with males of species B only. This could happen in some of our populations, where pebbles and stones are more or less covered over by sand from one season to another, and the density of individuals is low (e.g. Fig. S1 in supplementary material from Ribardière et al., 2017).

In conclusion, it is unlikely that sexual isolation has evolved in response to- or as a by-product of divergent ecological conditions, but unstable habitat conditions may currently modulate the efficiency of sexual isolation through stochastic variations in the density and distribution of individuals.

### Sexual isolation is not strict, and it is asymmetric

Sexual isolation was strong, but it was not 100%. Hence sexual isolation is not sufficient to prevent interspecific gene flow entirely on its own. As a result, when ecological isolation is lacking, then hybridization is expected to happen now and then.

Moreover, results from the free-choice experimental population showed sexual isolation to be asymmetric (Figs. 4 and 5). In this experiment, *J. albifrons* females mated exclusively with *J. albifrons* males while *J. praehirsuta* females mated with *J. praehirsuta* (*n*=5), *J. albifrons* (*n*=1) and F1 hybrid males (*n*=2). Accordingly, pair sexual isolation indices for heterospecific pairs were smaller (indicating stronger isolation) when the female involved was *J. albifrons* rather than *J. praehirsuta* (Table S1). Summarizing male sexual phenotypes using the first axis of a principal component analysis (Fig. 5), we found that the probability that a male mated with a *J. albifrons* female matched the density distribution of *J. albifrons* male sexual traits. By contrast, the probability that a male mated with a *J. praehirsuta* female was clearly less concordant to the density distribution of *J. praehirsuta* male sexual traits (Fig. 5B). This shows that females *J. praehirsuta* mated with our three categories of males (*J. praehirsuta*, F1 hybrids, and *J. albifrons*) because they are less selective that *J. albifrons* females with regards to male sexual traits (illustrative of “type d” preference function in Fig. 5 from Ryan & Rand, 1993).

This asymmetry is also confirmed by the no-choice experiments, where inter-specific crosses were systematically more successful with *J. praehirsuta* females (this is consistent in crosses within Brittany, Normandy, and inter-regions, data not shown). Over all interspecific crosses, 4 out of 36 were successful when the female was *J. albifrons*, while 10 out of 28 were successful when the female was *J. praehirsuta* (Fisher’s exact test p=0.036, Fig. 6). Interestingly, the same result was obtained by M. Solignac with no-choice experiments using individuals from a population from France (Brittany) crossed with individuals from Germany (0% success for German *J. albifrons* females crossed with French *J. praehirsuta* males vs 22 to 46% success for French *J. praehirsuta* females crossed with German J. *albifrons* males, depending on the method of calculation, Solignac, 1978).

**Figure 6.**
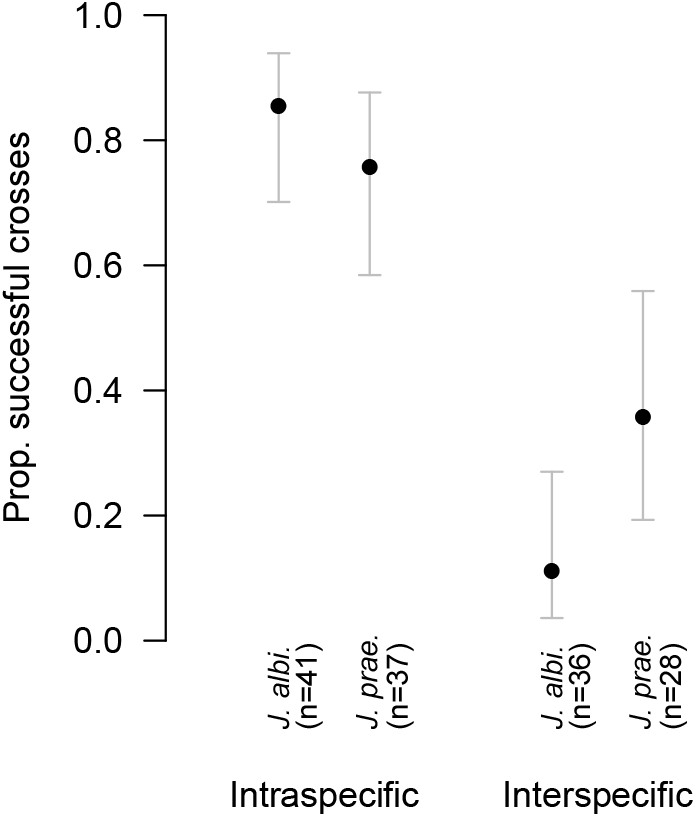
Proportion of successful intra- and interspecific crosses in no-choice experiments involving either *J. albifrons* or *J. praehirsuta* females. The sample sizes (number of experimental crosses) are given in brackets, and the bars give 95% confidence intervals around each observed proportion. A cross was successful if it produced at least one offspring. Sexual isolation between species is stronger in one direction (*J. albifrons* female).

The asymmetry in sexual isolation is thus observed in all conditions: large-scale allopatry (between countries, Solignac, 1978), smaller-scale allopatry (between regions, this study), sympatry with ecological isolation and no hybridization (Brittany, this study), and sympatry without ecological isolation and with introgressive hybridization (Normandy, this study). This is interesting because it suggests that the asymmetry has a general cause of ancestral origin and has not evolved in response to local conditions, in agreement with our previous conclusions that sexual isolation has not evolved following context-dependent sexual selection mechanisms. The forces that can potentially cause an asymmetry in sexual isolation (discussed e.g. in Coyne & Orr, 1998, Svensson et al., 2007) are tied to the evolution of sexual isolation itself, which we discuss now.

### Did sexual isolation between J. albifrons and J. praehirsuta evolve from sexual selection ?

We follow Coyne and Orr’s definition of sexual isolation (Coyne & Orr, 2004, p. 213 and Table 1.2 p. 28-29): it can include behavioural isolation and different elements of copulatory and gametic isolation. We know from previous work that behavioural isolation is the major component of sexual isolation in the *Jaera albifrons* group, and that there seems to be no copulatory (Bocquet, 1953, p. 297-298) or non-competitive gametic isolation (Solignac, 1978, p. 80-82). It is totally unknown, however, whether competitive gametic competition happen. We have shown in this study that females can store sperm from at least two males for several months (see results and supplementary information), which may set the stage for sperm competition and/or cryptic female choice. This aspect deserves further empirical investigation, and we discuss below only the evolution of the behavioural component of sexual isolation (see Coyne & Orr, 2004, p. 217 for a review of the forces that can cause the evolution of behavioral isolation).

Male sexual traits in *J. albifrons* and *J. praehirsuta* are used only for courtship behaviour and bring no other direct benefit to the bearer (e.g. advantage in local adaptation, Tinghitella et al., 2009, or in male-male competition, Sefc et al., 2015), and there are also no obvious direct benefits or costs for choosers (e.g. resource provided by mating partners, or, on the contrary, detrimental effects of mating, e.g. Svensson et al., 2007). The differences between males also involve clearly distinct, nonoverlapping characteristics that are not compatible with a loss of courtship elements in one of the two species (Kaneshiro, 1980, Ryan & Wagner, 1987). In addition, in this study we suggested that sexual isolation did not evolve alongside particular ecological conditions. These observations together eliminate many potential evolutionary forces, leaving models of sexual selection involving indirect benefits for female choice (e.g. Fisher-Lande process, or good- or compatible genes models) as the most plausible driver of sexual isolation between *J. albifrons* and *J. praehirsuta*. Under this hypothesis, sexual selection would also have driven the asymmetry in sexual isolation. This hypothesis is based on indirect deductions and will now need empirical testing (e.g. following Panhuis et al., 2001).

### Why do *J. albifrons* and *J. praehirsuta* coexist in hybridizing populations?

Individuals identified as *J. albifrons* in region Brittany (with ecological isolation) belong to the same biological species as individuals identified as *J. albifrons* in region Normandy (where there is no ecological isolation), and likewise for species *J. praehirsuta*. This was argued by taxonomists and evolutionary biologists based on the observation of secondary sexual traits (Bocquet, 1953, Prunus, 1968, Solignac, 1978) and confirmed in this study using cross experiments: conspecific crosses gave the same results within and across regions (no reproductive isolation between regions).

However, we also know that reproductive isolation is not complete in populations without ecological isolation, where hybridization leads to introgression (Bocquet & Solignac, 1969, Solignac, 1969a, 1978, Ribardière et al., 2017). This situation raises questions about the conditions of coexistence of the two species in spite of hybridization and ecological equivalence (e.g. Coyne & Orr, 1998). Ribardière et al. (2017) discussed the peculiar nature of the hybridizing populations, emphasizing that they seem to receive no influx of individuals from pure parental populations, and that no fine-scale ecological differentiation was observed (the two species are repeatedly found to be intermingled in a number of different sites). The authors suggested sexual and post-zygotic isolation as two potential forces somehow acting to keep the species isolated. Here we found sexual isolation to be effectively strong in hybridizing populations, but very little support for first-generation post-zygotic mechanisms (*AC*_2_ and *AC*_3_ added to −0.04 and −0.06 in regions Brittany and Normandy, Table 2). We could not quantify post-zygotic isolation from the free-choice experimental population because only one heterospecific mating pair and two backcrosses (F1-hybrid fathers) produced offspring. Yet these broods showed no sign of reduced fitness in any way (brood size and offspring survival, data not shown). Post-zygotic barrier effects need to be investigated further, but so far sexual processes (mate choice and/or intra-sex interactions) constitute the only force identified that could maintain the two species separated in spite of extensive introgression. This alone is unusual, and the fact that the two species are found to coexist in different places and on the long term despite ecological equivalence poses another conundrum that deserves further investigations.

### Conclusion & perspectives

In this study we looked at two closely related species that are reproductively isolated by ecological and sexual barriers, asking first if sexual isolation could maintain species integrity on its own or is only acting secondarily alongside ecological isolation, and second if sexual isolation may have evolved independently of local ecological context. We found that sexual isolation is a strong barrier that does not fall apart when ecological isolation is absent, thereby indicating that it cannot be ruled out as a primary driver of reproductive isolation. And we found that sexual isolation most probably evolved independently of local ecological conditions. These results suggest that the *J. albifrons* / *J. praehirsuta* pair is an example where sexual isolation evolved and acts largely independently of other forces (at the exception, perhaps, of local ecological disturbance that could modulate mate choice options through variations in density). Yet, sexual isolation alone falls short of ensuring total reproductive isolation, and ecological isolation seems to be necessary for reproductive isolation to be complete.

This pair of species shows many indirect clues suggesting that sexual isolation evolved directly from sexual selection, but whether or not it initiated speciation remains an open question. One way forward with this question will be to assess whether replicates of mixed populations occupy a range of different habitats. If sexual isolation did drive speciation, one predicts to find the *J. albifrons* / *J. praehirsuta* pair in different habitat settings (the strongest evidence would come from populations where the two species have reversed ecological niches compared with the ones that have been described so far). Given the extended geographic distribution of these two species (American and European coasts of the temperate and cold waters of the Northern-Atlantic Ocean), this prediction is testable. Finally, the conditions of coexistence of these two species in hybridizing populations are another open question. For that, further examination of post-zygotic barrier effects, fine-scale tests for habitat specialization (unseen so far), and interactions favouring density-dependence mechanisms (such as species-specific pathogen effects) may provide some answers.

## Acknowledgements

We thank Carole Smadja for discussions and insightful comments on this article, and Sébastien Collin for the confocal laser scanning microscope images used in figures 1 and 5. Martine Maan and an anonymous reviewer provided many useful comments on an earlier version of the manuscript. This work benefited from access to the Biogenouest genomic platform at Station Biologique de Roscoff and was funded by the French Agence Nationale de la Recherche (grant ANR-13-JSV7-0001-01 to T.B.).

## Author contributions

Conceptualization and methodology: AR and TB. Field sampling and species identification: AR, EP, JC, CH, SL and TB. Crossing experiments and maintenance of the individuals in the laboratory: AR, EP, JC, CH, SH, and TB. Genotyping and phenotyping: AR, EP, JC, CDT, CH and TB. Analyses and writing: AR, EP, and TB. Supervision, project administration and funding acquisition: TB.

## Conflict of interest disclosure

The authors of this preprint declare that they have no financial conflict of interest with the content of this article

## Data accessibility

All data are available from the corresponding author on request.

