## Supplementary material for "Sexual isolation with and without ecological isolation in marine isopods *J. albifrons* and *J. praehirsuta*"

**Supplementary information**

**1- Females sampled in natural populations**

All controlled crosses were set-up using individually reared, virgin individuals that were born in the lab from females previously sampled in natural populations. For intra-region crosses, these females were sampled between March 29 and April 20, 2015. For inter-region crosses, the females were sampled between the $14^{th}$ and $24^{th}$ of November, 2016. Sampling locations are indicated in main text figure 2.

**2- Rearing conditions of adult females and offspring**

All female adults (see preliminary steps in Fig. 1) were reared individually in 6-well plates (one individual per well, each well has a diameter of approximately 3.5 cm and contains ca. 10 mL of 3 µm-filtered seawater). Following the conditions described in Bocquet (1953, p. 212-213), each well contained small pieces of green algae (*Enteromorpha* sp.) and a small piece of elm leaf (elm leaves provide shelter and food, and have been used as standard conditions for several decades by previous researchers, e.g. Solignac, 1978). The plates were kept in thermostat cabinets at 17°C with an 11h/13h light/dark cycle and seawater was changed once a week (together with algae and leaves when needed). Embryos develop in a marsupium (brood pouch containing typically ca. 10-15 offspring) until the female releases them. At that stage the offspring measure approximately half a millimeter and closely resemble the adults (direct development, no pelagic larval stage).

Offspring were isolated one by one within a few days after their release and reared individually in the same conditions as the adults, with two exceptions: seawater was not changed during the first week, and elm leaves were added only during the second week. Under these conditions, individuals can be sexed around the age of 4 to 5 weeks.

**3- Conditions of no-choice experiments**

In no-choice experiments, controlled crosses were performed by pairing one male and one female within a unique well. These experiments were run in two batches. Intra-region / intra-specific crosses (n=46) and intra-region / inter-specific crosses (n=34) were set-up in June 2015, while inter-region crosses (40 intra-specific and 40 inter-specific crosses featuring individuals from opposite regions) were set-up in January 2017.

In order to avoid manipulation errors, we used only two wells per plate (as represented in Fig. 2). Each cross was checked for offspring at least two times per week until offspring were produced or one of the parents died (up to 196 days). All offspring were reared in the conditions described above. Many pairs were kept long after they produced a first brood, thereby allowing us to collect and rear successive broods, but all the data analysed in this article refer to the first brood only.

**4- Detailed protocol for the free-choice experiment**

A mixture of 30 virgin females and 51 virgin males of controlled origin composed our experimental population (Fig. 2). These individuals are the "parents", themselves produced in the lab (Fig. 2) from known mothers and fathers that we call "grandparents".

All parents were photographed individually using a binocular microscope and a 10mm graduated scale for later size measurements in ImageJ v.2.0.0. Additional close-up pictures of the head of individuals (featuring recognizable pigmentation patterns) were also taken for downstream identification once the individuals will be retrieved (in particular for female identification, see below).

All adults were put together in a small aquarium (26 × 17 cm, 13 cm deep) on November 19th, 2015. The aquarium was maintained at 17°C using a thermostated water bath, and contained the same oxygen and food sources used to raise the individuals since their birth in the lab. These conditions were maintained for 12 days, during which all individuals could interact freely. This is the minimum time required for a female to produce offspring if such a female would have been fertilized early in the experiment (Solignac, 1976). Four individuals were found dead during this 12 days period (see Results). All surviving females (n=27) were then removed from the aquarium and placed individually in 6-well plates (standardized conditions as in paragraph 2 above) that were then checked for offspring 2 to 3 times per week for 66 days.

The first offspring were observed on December 3rd (i.e. 3 days after females were isolated), and this part of the experiment was terminated on February 4th (i.e. 66 days after isolation) when a single female was still alive and had not produced any offspring. During these 66 days all the offspring produced were isolated in 6-well plates and reared individually (see conditions in paragraph 2 above). If a mother's first brood contained more than 10 individuals then the mother was photographed and fixed in ethanol. Otherwise it was kept alive in the same conditions until it produced a second brood in order to enhance downstream parentage assignment (see below). As it turned out, five females were allowed to produce two successive broods, but in all cases these females had a unique, identical mate across their two broods. For consistency, all downstream analyses were based on data from each female's first brood only (this is important e.g. for estimating male and female total reproductive output).

All offspring were reared for at least 57 days (up to 70 days) during which their survival was checked once per week. They were then sexed, photographed, and fixed in ethanol. In addition, the secondary sexual traits of all male offspring (Fig. 1) were examined using a microscope. For each male we recorded the presence and approximate size class (small vs large) of a carpal lobe on peraeopods 6 and 7 (P6 and P7), the number of spines, straight setae, and curved setae on the carpus of P6 and P7, the number of distal setae on P6 and P7, and the number of curved setae on the propod, carpus, and merus of peraeopods P1 to P5. It was not always possible to count curved setae precisely when there were more than 25-30 setae on a single peraeopod segment. In these cases the number of setae was approximated. Whenever this was possible, the same actions (sexing, measurements, phenotype description, and fixation) were carried for the offspring that were found dead in the course of the experiment.

Males of the parental generation were not removed right after the females were isolated because at that stage we did not know whether the 12-days period of free interaction had been sufficient for a large sample of females to be fertilized. Hence we kept the males for 20 to 24 additional days in the original aquarium, and the surviving males (n= 43) were then photographed, their phenotype was described (using the same traits as described above for male offspring), and they were fixed in ethanol. When possible, all such males that were found dead before that time were processed similarly. Male phenotypes were summarized using principal component analyses (PCA) based upon the following 13 phenotypic variables: carpal lobe on peraeopods 6 and 7 (P6 and P7) coded as 0 (absent), 0.5 (small), or 1 (large), number of spines on P6 and P7, total number of straight or curved setae on the carpus of P6 and P7, number of distal setae on P6 and P7, and total number of curved setae on peraeopods P1 to P5 (i.e. adding setae counted on each segment). The PCAs were run in R v.3.3.3 (R Core Team, 2017) using package ade4 v.1.7.6 (Chessel et al., 2004).

**5 - Parentage analyses and female identification in the free-choice experiment**

We used genetic parentage analyses to achieve two goals. First, while the females that entered the experimental population were identified as *J. albifrons* or *J. praehirsuta* based upon the type of controlled cross from which they originated, they could no longer be identified once they were mixed up altogether and later isolated one by one until they produced offspring (because the females of each species are nearly morphologically identical). Hence we used DNA data from the individuals that produced these females (the "grandparents" used in controlled crosses previously performed in the lab) in order to re-identify each female through genetic parentage assignment. Female identification was aided by photographs taken before and after the experiment (see below). Second, we used parentage analyses to identify the fathers that produced each offspring (the mothers were known since females were kept individually until they produced offspring, Fig. 2). To achieve these goals, all offspring, potential parents, and grandparents were genotyped at 13 microsatellite loci (Ja02, Ja13, Ja21, Ja22, Ja23, Ja30, Ja35, Ja39, Ja41, Ja64, Ja71, Ja78, and Ja94, Ribardière et al., 2015). DNA extraction, PCR amplification, and allele scoring were performed following Ribardière et al. (2015).

Multilocus genotypes were then used for parentage assignment in the software Colony v2.0.6.1 (Jones & Wang, 2010). For female parentage assignment we set options for male and female polygamy, long run of the full likelihood method with high likelihood precision. While the primary goal of this analysis was only to sort *J. albifrons* vs *J. praehirsuta* females, it further allowed us to determine the best parameter options because we could check parentage results against 1) the potential pairs of parents that produced the females and males mixed in the experimental population (these potential pairs were known because they corresponded to our controlled crosses) and 2) photographs of each male and female taken before and after the experiment. Combining results from the parentage analyses with photograph identification of all parents allowed us to achieve a reliable identification of each parent (that is, we did not only re-classify each female parent as being a *J. albifrons* or *J. praehirsuta*, but we also re-identified each male or female parent so that we could know exactly who it was and from what controlled cross it was obtained initially).

The second objective of parentage analyses was to assign all offspring produced in the experimental population. The mothers of these offspring were known, since all females had been kept isolated in six-well plates until they produced juveniles. Here the objective was to find what father sired each offspring. For that we used the same parameter options, with the addition of known maternal sibship information in Colony.

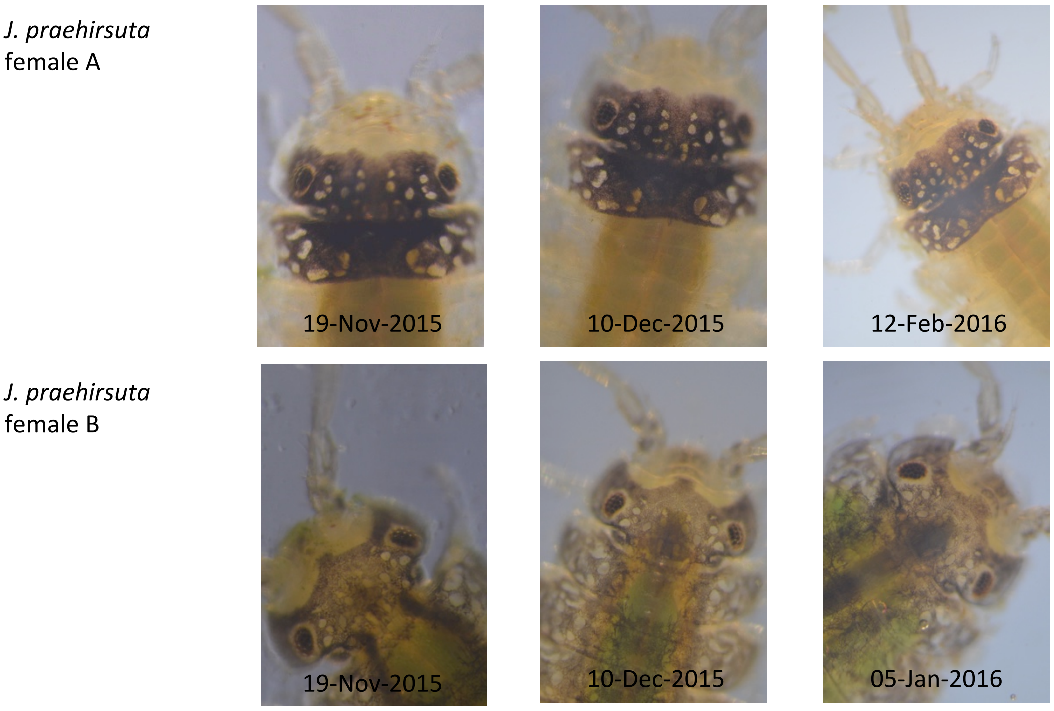


**Figure:** Examples of close-up pictures taken using a reflex camera mounted on a binocular microscope. These pictures show two *J. praehirsuta* female parents that were photographed before the free-choice experiment (Nov. 19, 2015), once during the experiment (Dec. 10, 2015), and finally when the females were fixed in ethanol for genetic analyses (early 2016). Not all individuals are so obviously different, but most can be re-identified easily using general pigmentation patterns (more or less extended black pigments) and details of white spots on the head of individuals. Here we used pictures as a control in addition to parentage analyses to primarily assess what females belonged to *J. praehirsuta* vs *J. albifrons*, but the combination of parentage results and photographs allowed us to go further and actually identify each individual (males and females) one by one. Variations in pigmentation patterns are one reason that motivated some early work on the *Jaera albifrons* complex, and were used before as a marker to identify individuals in mating experiments (Solignac, 1978).

### 6- Analysis of raw data for the free-choice experiment

Out of 30 potential parent females, 7 died during the experiment (no offspring produced), 22 produced offspring (14 *J. albifrons* and 8 *J. praehirsuta*), and the last one (*J. praehirsuta*) was watched for 66 days (once pulled out from the experimental population) but did not produce any offspring.

Out of 51 males, 1 male was found dead one day before females were removed from the experimental population, and 2 were never found (and thus could have died early in the experiment without being detected). But 2 out of these 3 males actually did sire offspring. So it leaves a maximum of 1 male out of 51 for which we don't know if it died too early to have a chance to court females. We considered that this uncertainty was negligible (that is, all further computations and interpretations are based as if all males had equal chances to reproduce).

More importantly, the parentage analyses revealed two errors in the experimental setup. First, a set of four brothers that were thought to originate from an interspecific controlled cross ("grandparents": female *J. praehirsuta* / male *J. albifrons*) in fact did not originate from this cross. Before being taken out and used in the controlled cross, their mother had mated with one of her brothers that managed to crawl between wells in the 6-well plate where this family was raised (see preliminary steps in Fig. 2). Fortunately, these four males (inbred *J. praehirsuta* instead of "hybrids") did not sire any offspring in the experimental population and they were simply removed from all downstream analyses. The second error is similar: the very first female that produced offspring after being taken out of the experimental population in fact had been mated by one of her brothers before the experiment. Hence this female and her brood were also taken out from all downstream analyses. No other issues were detected: parentage analyses showed that all other parents corresponded to our controlled crosses and all other offspring were conceived in the experimental population.

In summary, all downstream analyses were based on reproduction patterns for 47 males and 22 females. These individuals produced 359 offspring, out of which 239 could be genotyped (that is, offspring that survived for 57-70 days until they were genotyped or were found dead in the course of the experiment but could be correctly genotyped). Overall, 5 females were kept alive until they produced a second brood (3 *J. praehirsuta* and 2 *J. albifrons* females, 1 to 17 offspring per brood). Each of these second broods were parented by the same single male as the first broods, so that these second broods did not affect our estimates of mating success. These second broods were removed from all downstream analyses, and reproductive success measures were finally based on 218 offspring stemming from the first brood of each female parent.

**7- Fecundity, polyandry, and multiple-paternity**

*J. albifrons* females produced from 3 to 31 offspring per brood (*n*=14, mean=16.64), while the fecundity of *J. praehirsuta* females seemed slightly lower (5 to 19 offspring per brood, *n*= 7, mean=11.71, excluding the one female that produced no offspring at all), although this difference was not significant (*n*=21, generalized linear model *p*=0.146). Female fecundity was significantly related with body size (*n*=18 females for which we had body size at the time when the first brood was released, generalized linear model *p*=0.022). There is little power to test this, but the three *J. praehirsuta* females that mated with heterospecific males did not show any sign of reduced fecundity (brood size 7, 15, and 19 offspring, and the female that released only 7 offspring in her first brood produced a second brood of 17 with the same heterospecific father).

As mentioned above, 5 females had 2 male partners (4 out of 14 reproductive *J. albifrons*, and 1 out of 7 reproductive *J. praehirsuta*). The four *J. albifrons* females mated with two distinct *J. albifrons* males, while the *J. praehirsuta* female mated with one *J. praehirsuta* and one *J. albifrons* male. All 5 cases corresponded to multiple paternity within the first brood (and not to two different fathers in two different broods). These broods contained 15 to 23 offspring (7 to 10 genotyped offspring), and the minimum contribution of a given father was 0.14.

**Supplementary tables and figures**

Table S1 - Pair sexual isolation (PSI) between two types of females (*albifrons* and *praehirsuta*) and three types of males (*albifrons*, *praehirsuta*, and hybrids) in the free-choice experiment. Sample size (number of individuals and number of successful pairs of each type) are indicated in brackets. PSI values below 1 indicate a deficit of observed pairs, while values above 1 indicate an excess of observed pairs (Rolan-Alvarez & Caballero, 2000).

| | Males | | |
|---|---|---|---|
| Females | *J. albifrons* (17) | hybrid (13) | *J. praehirsuta* (17) |
| *J. albifrons* (14) | 1.37 (n=18) | 0 (n=0) | 0 (n=0) |
| *J. praehirsuta* (8) | 0.17 (n=1) | 3.25 (n=2) | 3.25 (n=5) |

Figure S1

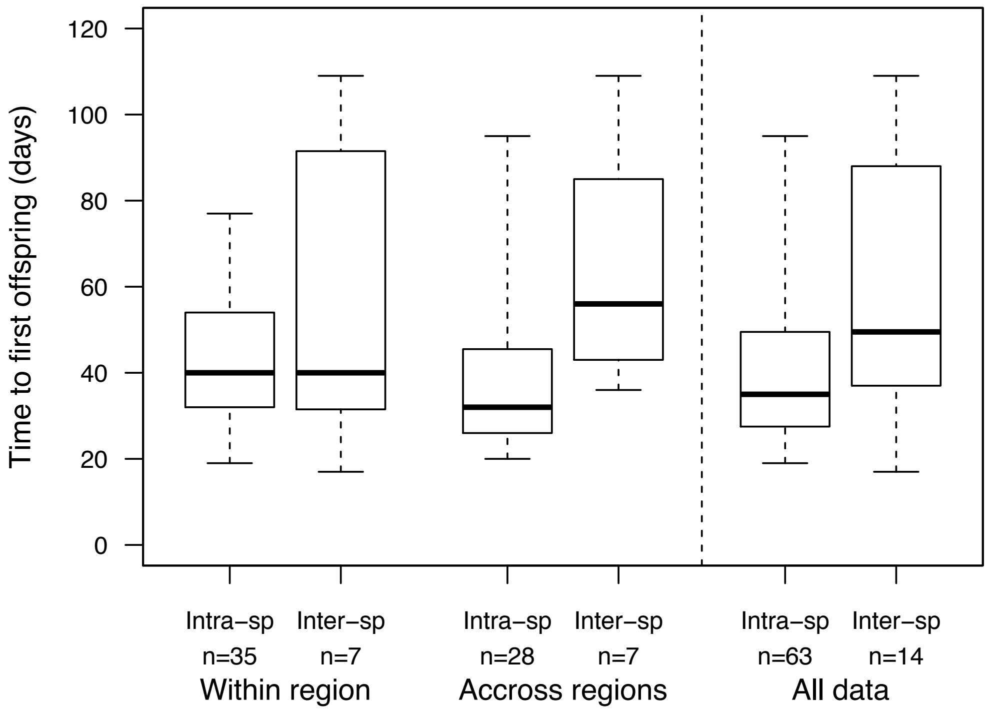


Figure S1 - Delay between the start of each cross (when a virgin male and a virgin female are put together) and the release of the first brood. It takes longer for intraspecific pairs to produce offspring, indicating sexual isolation between species.

Figure S2

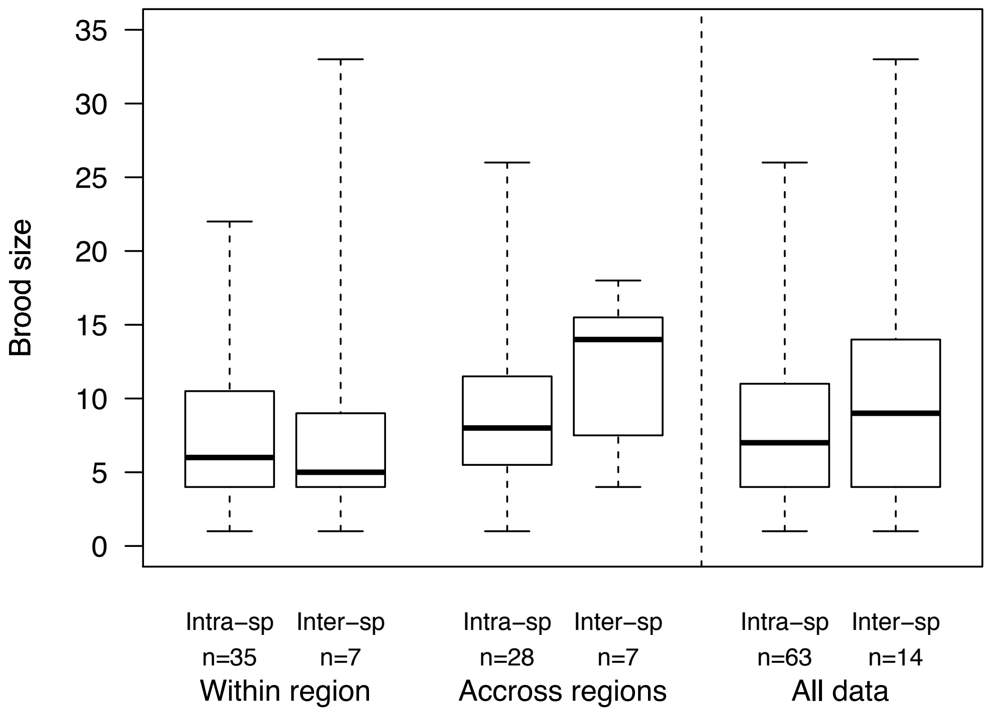


Figure S2 - Number of offspring released per brood obtained from experimental no-choice intraspecific crosses (either *Jaera albifrons* or *J. praehirsuta*) and inter-specific crosses. The sample size (number of broods) is given below each box. There is no difference in brood size between cross types, indicating that no post-zygotic barrier is acting at this stage (embryo development in F1 hybrids).

Figure S3

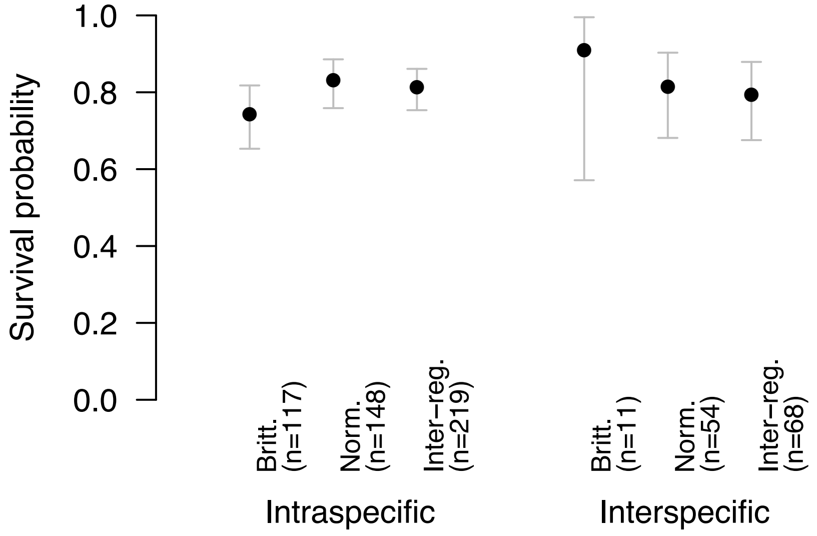


Figure S3 - Proportion of offspring surviving at day 35. Offspring stem from experimental no-choice intraspecific crosses (either *Jaera albifrons* or *J. praehirsuta*) and inter-specific crosses. The male and female of a given cross could come from the same region (Brittany or Normandy, see text) or each from a different region (inter-reg. crosses). The sample sizes (number of experimental crosses) are given in brackets, and the bars give 95% confidence intervals around each observed proportion of offspring surviving.

Figure S4

A

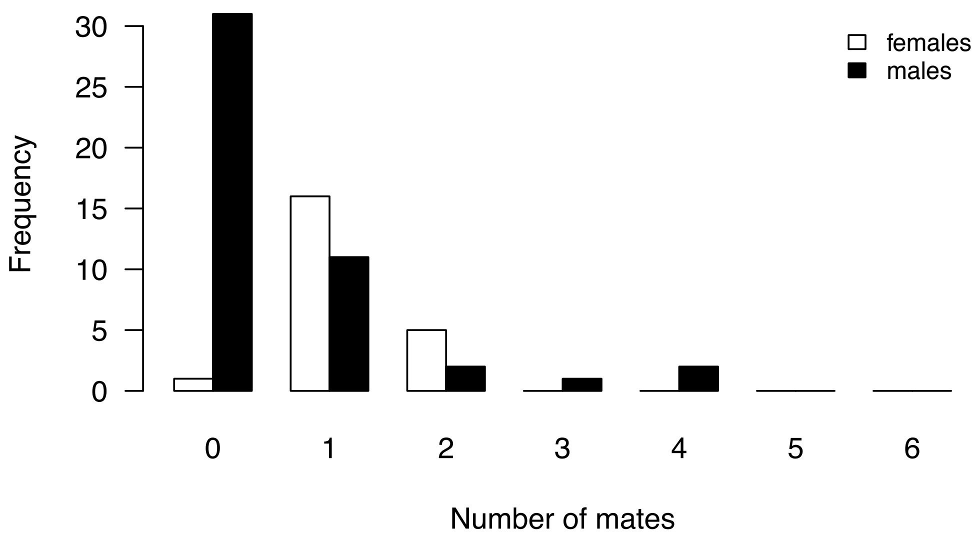

females
males
Frequency
30
25
20
15
10
5
0
0
1
2
3
4
5
6
Number of mates


B

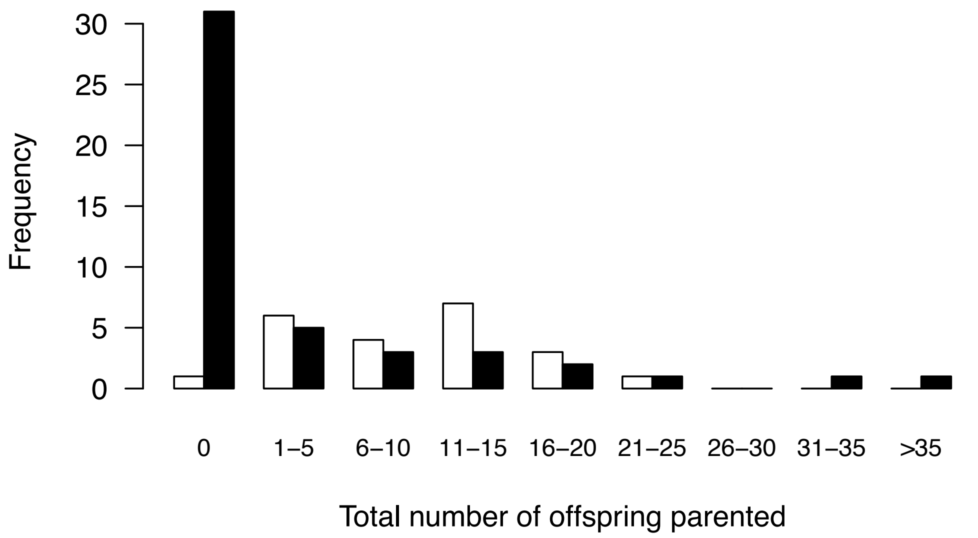

Frequency
30
25
20
15
10
5
0
0
1–5
6–10
11–15
16–20
21–25
26–30
31–35
>35
Total number of offspring parented

Figure S4

Distribution of male and female mating success (A) and total reproductive success (B) in our experimental free-choice population (regardless of species or phenotype). Female data are shown in white, male data in black. The total reproductive success (panel B) is defined as the number of offspring genetically assigned to a given adult male or female. That is, it does not include the offspring that were known to be released by a given female but did not survive up to the genotyping step of the protocol described in figure 1. This figure shows the difference between male and female variance in breeding success.

Figure S5

A

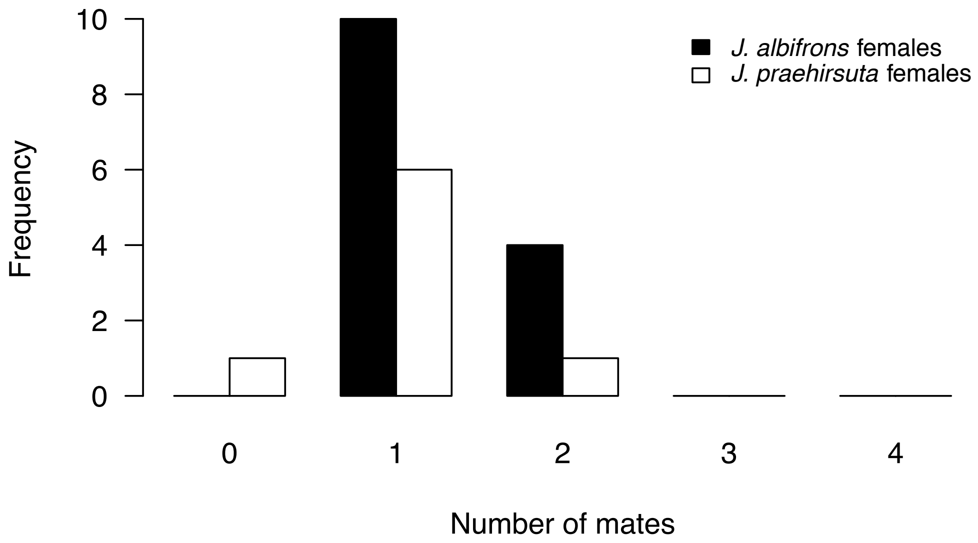


B

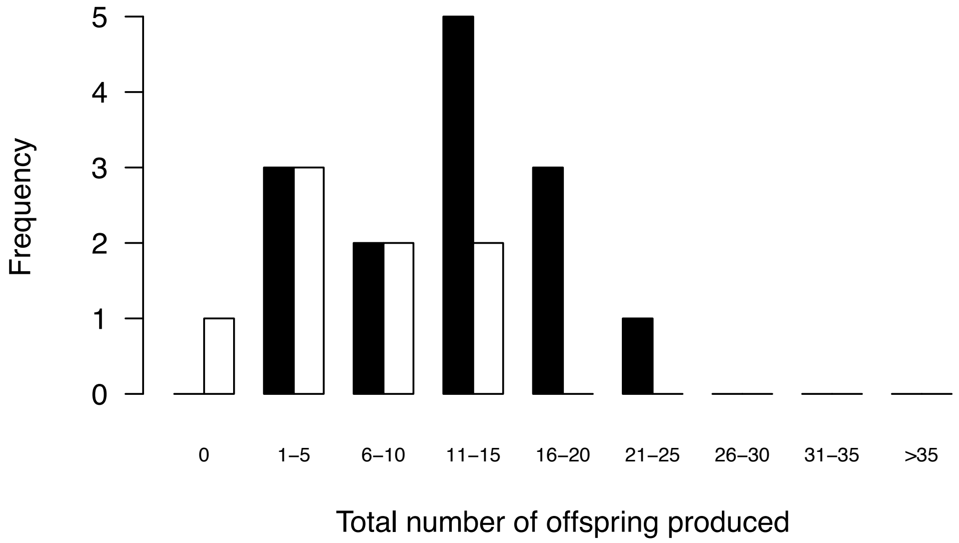


Figure S5

Distribution of female mating success (A) and female total reproductive success (B) for species *Jaera albifrons* and *J. praehirsuta* in the free-choice experiment. In this figure we see that mating success and total reproductive success were higher for *J. albifrons* females than *J. praehirsuta* females.

Figure S6

A

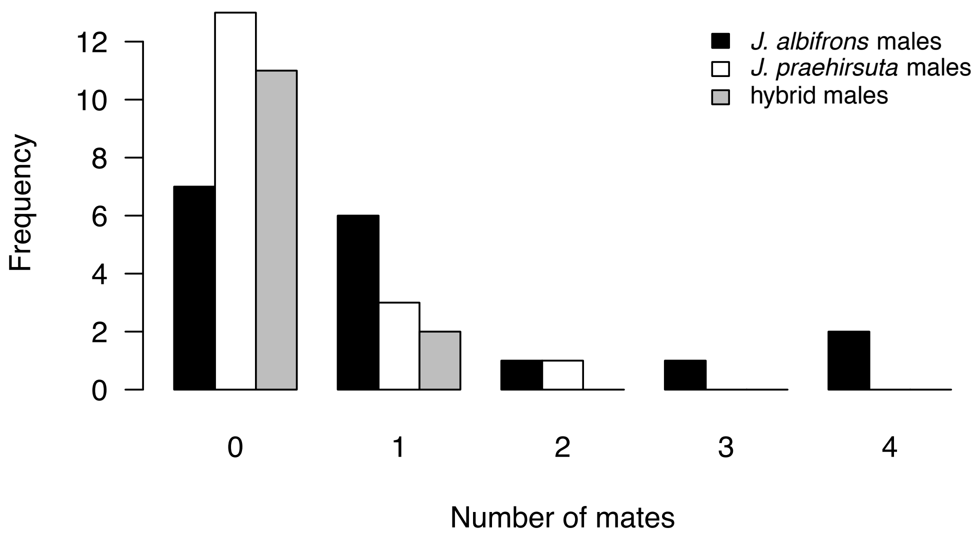


B

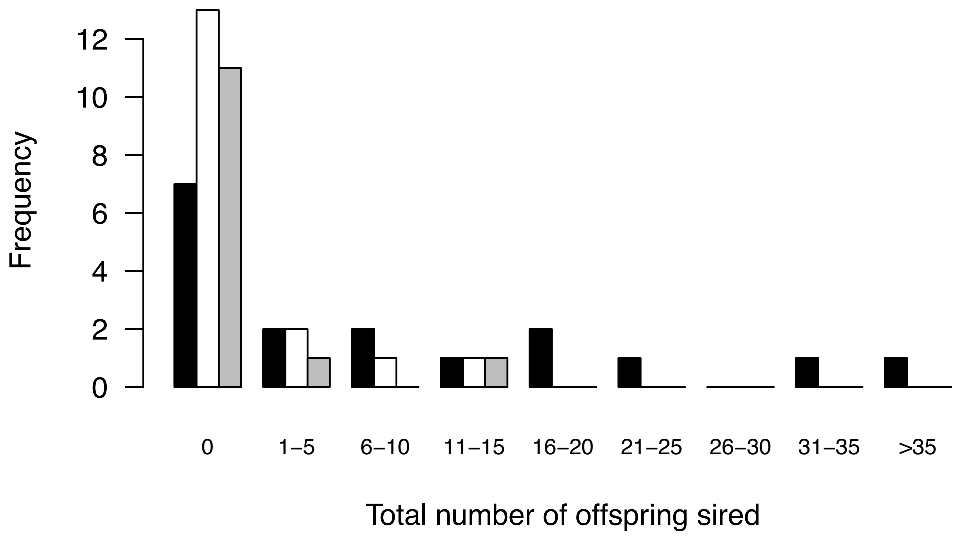


Figure S6

Distribution of male mating success (A) and male total reproductive success (B) for each male type (*albifrons*, *praehirsuta*, and hybrids) in the free-choice experiment. In this figure we see that mating success and total reproductive success were highest for *albifrons* males, intermediate for *praehirsuta* males, and lowest for hybrids.
